# Highly pathogenic avian influenza management in high-density poultry farming areas

**DOI:** 10.1101/2025.03.25.645233

**Authors:** Claire Guinat, Cecilia Valenzuela Agüí, François-Xavier Briand, Debapriyo Chakraborty, Lisa Fourtune, Sébastien Lambert, Andrea Jimenez Pellicer, Severine Rautureau, Guillaume Gerbier, Louis du Plessis, Tanja Stadler, Beatrice Grasland, Mathilde C. Paul, Timothée Vergne

**Affiliations:** Interactions Hôtes-Agents Pathogènes (IHAP), Université de Toulouse, INRAE, ENVT, Toulouse, France; Department of Biosystems Science and Engineering, ETH Zurich, Schanzenstrasse 44, 4056 Basel, Switzerland; Swiss Institute of Bioinformatics (SIB), Lausanne, Switzerland; French Agency for Food, Environmental and Occupational Health & Safety (ANSES) Laboratory of Ploufragan-Plouzané-Niort, Ploufragan, France; Geomatys SAS, 1000 Avenue Agropolis, Montpellier 34394, France; Direction Générale de l’Alimentation (DGAL), Ministère de l’Agriculture et la Souveraineté Alimentaire, Paris, France

**Author notes:** Corresponding author: Claire Guinat. These authors contributed equally (Cecilia Valenzuela Agüí, François-Xavier Briand).

## Abstract

The continuous spread of highly pathogenic avian influenza H5 viruses poses significant challenges, particularly in regions with high poultry farm densities where conventional control measures are less effective. Using phylogeographic and phylodynamic tools, we analysed virus spread in Southwestern France in 2020-21, a region with recurrent outbreaks. Following a single introduction, the virus spread regionally, mostly affecting duck farms, peaking in mid-December with a velocity of 27.8 km/week and an effective reproduction number between farms (*R*_e_) of 3.8, suggesting the virus can spread beyond current control radii. Transmission declined after late December following preventive culling. Farm infectiousness was estimated around 9 days. Duck farm density was the main driver of virus spread and we identified farm density and proximity thresholds required to maintain effective control (*R*_e_ < 1). These findings offer actionable guidance to support regional biosecurity and to improve the robustness of the poultry sector to mitigate future outbreaks.

## Introduction

In recent years, the unprecedented and widespread outbreaks of highly pathogenic avian influenza (HPAI) H5 viruses, particularly clade 2.3.4.4b, have resulted in catastrophic socioeconomic and ecological impacts. Initially emerging in Asia, these viruses have been carried across continents by migratory wild birds, leading to devastating poultry outbreaks since 2016 ^1^. Alarmingly, since 2021, these viruses have shown endemic circulation in wild bird populations and recently expanded their host range to affect wild and domestic mammals not typically susceptible ^2,33/25/2025 7:18:00AM^. Moreover, sporadic human cases have been confirmed in multiple countries across four continents from 2020 to 2024 ^4^. The virus’ broad host and geographic ranges have heightened concerns about its potential to adapt to mammals, fueling fears of an HPAI pandemic in humans^5^. Consequently, managing HPAI outbreaks effectively has now become a critical global health objective ^6^.

France has been one of the most heavily affected countries in Europe by recurrent waves of HPAI since 2016 ^7–9^. In response to these waves, France has progressively implemented a series of regulations aimed at preventing introduction and spread of the virus. These included preventive culling, poultry movement restrictions, mandatory biosecurity training for poultry producers and biosecurity audits. In October 2021, the establishment of a high-risk diffusion zone (ZRD) refined the area of priority where measures had to be reinforced during high-risk periods such as duck flock pre-movement testing and compulsory indoor housing. Despite these efforts, outbreaks have continued to occur in regions with very high poultry farm densities, such as southwestern and western France, where the close proximity of farms likely makes these measures less effective. In October 2023, France initiated a vaccination campaign, showing promising results with fewer outbreaks compared to previous years. However, relying solely on vaccination may not be sustainable in the long term due to high costs, logistical challenges, mandatory surveillance protocols and the risk of trade restrictions ^10^. These challenges underscore the need to rethink the spatial distribution and organisation of poultry farms, particularly these high-density areas where tailored strategies are essential for effectively containing the virus and preventing its spread between farms ^11^.

Evaluating strategies for effectively stopping the spread of HPAI from one farm to another requires accurately reconstructing the epidemiological links between farms and identifying associated risk factors. This can be achieved by analyzing viral genetic sequences within a phylodynamic framework, which allows us to infer where and when the virus spread. By integrating generalized linear models (GLMs) into phylodynamic analyses, we can further identify the key drivers of virus spread ^12,13^. Using this joint approach, recent studies have identified significant associations of a range of factors, such as geographical proximity ^14–16^, poultry trade ^17–19^ and poultry population density ^20^, with virus spread in poultry. However, these studies were often limited by the number of available genetic sequences and the lack of precise location data, typically identified only at the country or regional level. As a result, past analyses were typically restricted to broad scales, making it difficult to draw actionable recommendations or develop effective control measures that account for local variations in farming practices or ecological conditions.

In this study, we build on these approaches to identify the key factors driving the spread of HPAI H5N8 viruses (clade 2.3.4.4b) between farms in Southwestern France in 2020-21 and provide options to enhance HPAI management in this high poultry farm density area. We applied a suite of phylogeographic and phylodynamic models to a comprehensive dataset of viral genetic sequences. Our phylodynamic analysis was extended with a GLM to account for a set of detailed variables being potential predictors of between-farm transmission such as poultry farm production and environmental data. Furthermore, our study leverages another dataset that distinguishes between operational and empty poultry farms during the study period. Our findings highlight the critical role that farm density and proximity on HPAI spread and derive practical recommendations to improve the robustness of poultry production sector against HPAI.

## Results

### Identifying the number of HPAI introductions to poultry farms

We analysed 432 genetic sequences of HPAI H5N8 viruses (clade 2.3.4.4b) from 468 confirmed infected poultry farms between December 2, 2020 and March 20, 2021. After subsampling to remove identical sequences with the same sampling date, species and location, our final dataset comprised 381 sequences, representing 83.1% of confirmed infected poultry farms (**Figure 1A**). As for the outbreaks, most sequences originated from infected duck farms, with a peak in early January 2021 (**Figure 1A**) and were from Southwestern France (**Figure 1B**). When combined with background sequences from Europe, the French dataset formed a monophyletic clade (branch support = 78.2%) (**Figure 1C**). The low genetic distance observed between the sequences and the fact they form a monophyletic clade suggest the introduction of a single viral lineage into the region, likely via one or more wild birds carrying genetically similar viruses within a short time frame, followed by sustained between-farm transmission.

**Figure 1.**
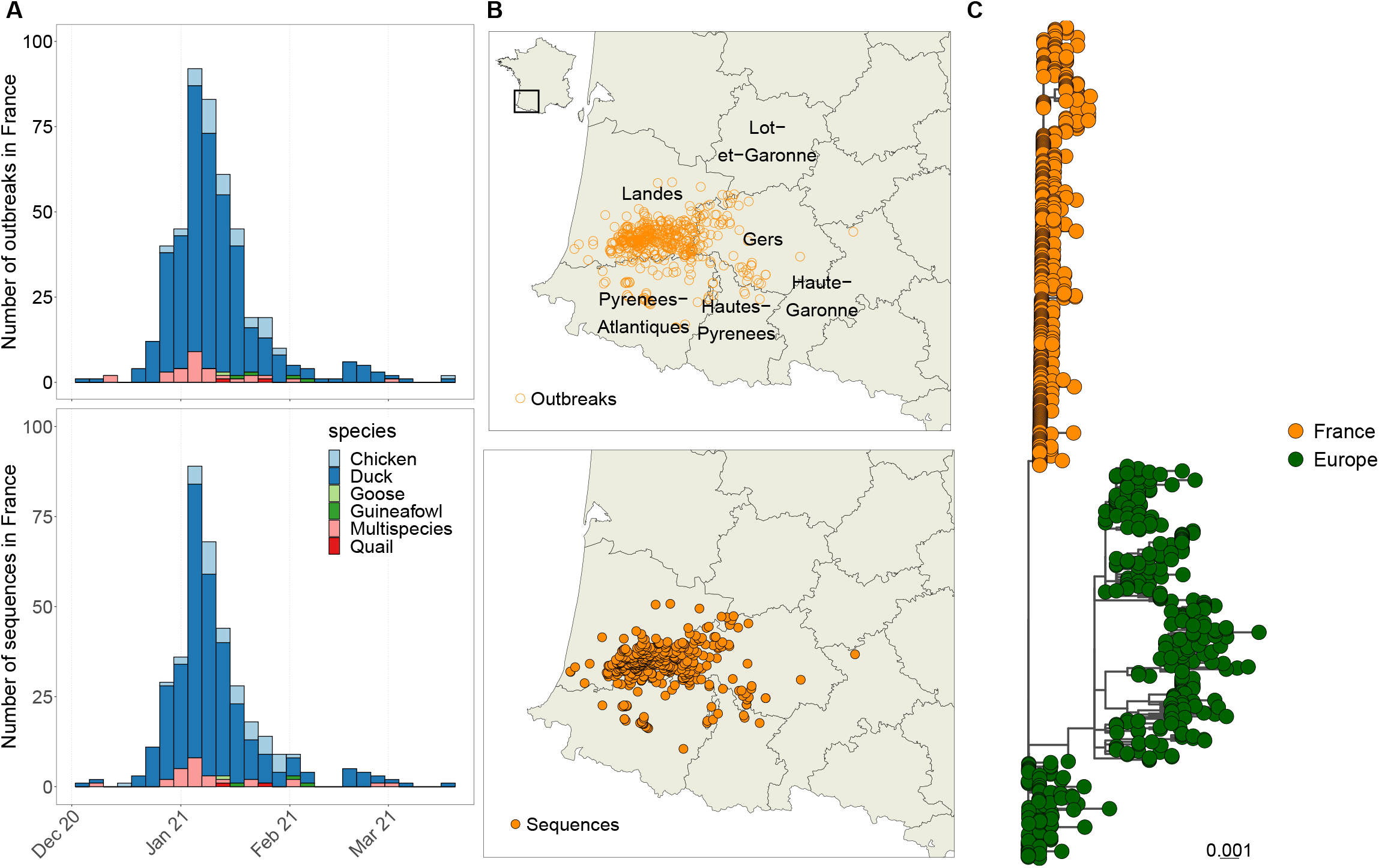
Distribution and phylogeny of HPAI H5N8 clade 2.3.4.4b genetic sequences from outbreaks in Southwestern France, 2020-21. The left and middle panel show the temporal (A) and spatial (B) distributions of HPAI H5N8 clade 2.3.4.4b outbreaks (n = 464) and genetic sequences (n = 381) in France in 2020-21. The right panel (C) shows the maximum likelihood phylogeny of the French genetic sequences (n = 381) along with background sequences from Europe from September 7, 2020 to May 1, 2021 (n = 342).

### Overview of spatio-temporal dispersal of HPAI

We investigated the spatio-temporal spread of HPAI between farms based on genetic sequences from the French dataset using a Bayesian continuous phylogeographic analysis.

Overall, the virus was maintained in duck flocks in Southwestern France with sporadic spillover events to other poultry species, in particular chicken flocks (**Figure 2A**). The virus initially emerged in Landes department, where it persisted from December 2020 to mid-February 2021, with subsequent transmission events to the southern Hautes-Pyrénées and Pyrénées-Atlantiques departments (**Figure 2B**). By mid-February, the outbreaks were largely under control, although a few local transmission events continued within the Gers department. We also estimated how fast HPAI spread between farms by inferring the weighted lineage dispersal velocity, which corresponds to the sum of the distance travelled along the phylogeny branch divided by the sum of the branch duration (**Figure 2C**). We found significant temporal variation in HPAI spread velocity, with an overall mean of 10.5 km/week (95% high posterior density interval, HPD: 6.1 – 16.5). The velocity peaked in mid-December 2020, reaching a maximum of 27.8 km/week (95% HPD: 14.2 – 49.5). After this peak, the velocity decreased in 2021 to an average of 9.5 km/week (95% HPD: 1.4-12.3).

**Figure 2.**
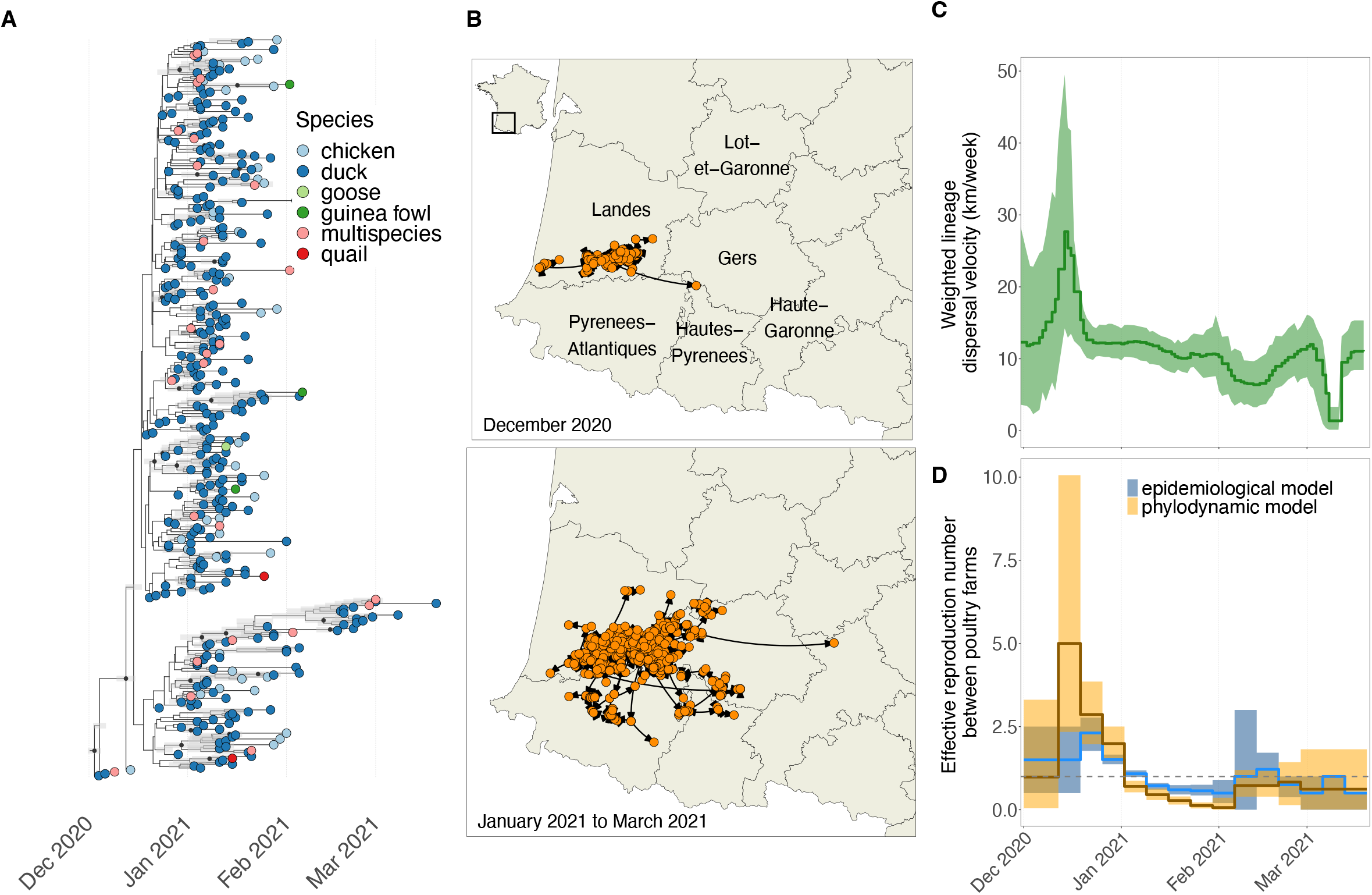
Transmission dynamics of HPAI H5N8 clade 2.3.4.4b between farms in Southwestern France, 2020-21. The left panel (A) shows the maximum clade credibility (MCC) tree of HPAI H5N8 virus sequenced from poultry farms during the 2020-21 wave in France. The tip colors represent the host species present in the infected farms. Grey bars represent the 95% highest posterior density (HPD) intervals reflecting uncertainty about internal node dates. The middle panel (B) represents the same MCC tree on a spatial map. The top right panel (C) shows the weighted lineage dispersal velocity over time with bold line representing the median values and the filled area represents the 95% highest posterior density intervals. The bottom right panel (D) shows the weekly estimates of the effective reproduction number (*R*_e_) over time inferred with a phylodynamic and epidemiological models. The bold line represents the median values and the filled area represents the 95% highest posterior density intervals and the 95% equal-tailed credible intervals. The horizontal black dashed line indicates a *R*_e_ value of 1.

### Estimates of HPAI transmission dynamics between farms

We inferred epidemiological parameters based on genetic sequences from the French dataset using a GLM extended birth-death-sampling model (phylodynamic model). The infectious period of poultry farms (defined as the time between infection and suspicion dates) was estimated at a median of 9.1 days (95% HPD: 7.8-10.5). The estimated effective reproduction number (*R*_e_) between farms (defined as the average number of infected farms infected by one infectious farm) over the study period showed significant temporal variation (**Figure 2D**). Generally, the peak *R*_e_ occurred in mid-December 2020, with *R*_e_ values significantly above 1 (median: 3.8; 95% HPD: 2.0 – 7.7), indicating heightened transmission risk during this period. Following this peak, *R*_e_ constantly decreased until end-January 2021 and reached values significantly below 1 during the month of January 2021. From early February to March 2021, *R*_e_ showed some resurgence in transmission, with values settled around 1. In addition to the phylodynamic approach, we estimated *R*_e_ based on epidemiological data only (epidemiological model) (**Figure 2D**). The pattern generally aligns with the trend observed with the phylodynamic model, although small differences were noted in the magnitude and precision of the *R*_e_ estimates.

### Predictors associated with HPAI transmission between farms

We assessed the contribution of predictors on the weekly *R*_e_ between farms based on genetic sequences from the French dataset using a GLM extended birth-death-sampling model. The inclusion probabilities and corresponding Bayes factors (BF) were used to evaluate the strength of each predictor’s contribution to the model. Two predictors, the number of active duck farms and active chicken farms (within a 10 km radius of an outbreak in the 7 days before the outbreak was reported) showed a decisive contribution (BF > 100) (**Figure 3A**). Note that *active* refers to farms with poultry present (excluding those that are empty due to sanitary procedures between production cycles, or due to culling measures, if they become infected or preventively culled). The density of active duck farms was positively associated with between-farm transmission, while the density of active chicken farms was negatively associated. The mean distance between active poultry farms and the percentage of water surface areas (within a 10 km radius of an outbreak in the 7 days before the outbreak was reported) indicated a strong contribution (BF > 10), with percentages of water surface areas positively associated with between-farm transmission and distances between active poultry farms negatively associated with lower between-farm transmission. Note that *poultry* refers to either ducks or chickens. The mean rainfall (7 days before the outbreak was reported) was not found as a significant predictor. Similar patterns were obtained when defining the predictors with a 12-day period instead of a 7-day period (**Figure S1**).

**Figure 3.**
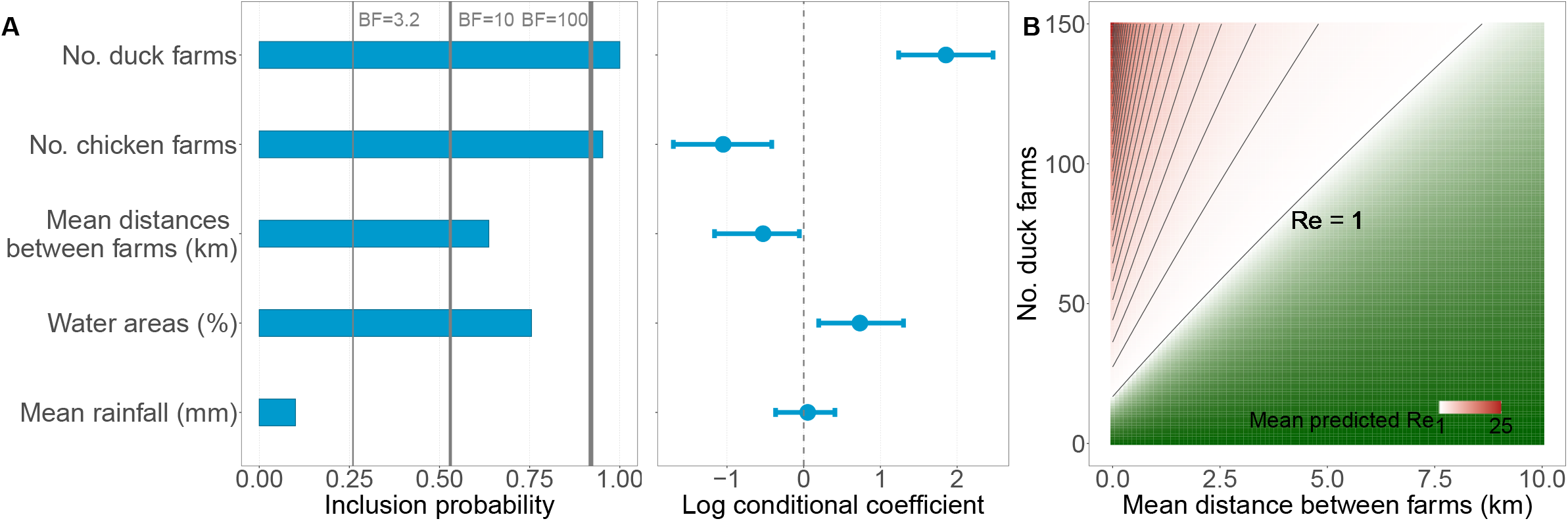
Drivers of HPAI H5N8 clade 2.3.4.4b transmission between farms in Southwestern France, 2020-21. The left and middle panel (A) shows the inclusion probability of the tested predictors for the weekly estimates of the effective reproductive number (*R*_e_) from the extended GLM birth-death-sampling model. This probability represents the proportion of the posterior samples in which each predictor was included in the model. Bayes Factors (BF) were used to determine the contribution of each predictor in the generalized linear model (GLM). BF quantify the likelihood of the posterior inclusion probability compared to the prior inclusion probability for each predictor. A cutoff of 3.2, 10 and 100 was used to indicate substantial, strong and decisive contribution of a predictor in the GLM, respectively ^54^. It also shows the log conditional median coefficients (points) and 95% highest posterior density interval (horizontal bars) for the predictors of *R*_e_, representing the (log) contribution of each predictor when it was included in the model. The right panel (B) shows the farm density and proximity thresholds to mitigate HPAI H5N8 clade 2.3.4.4b transmission between poultry farms in Southwestern France, 2020-21, with the predicted high 95% HPD *R*_e_ values as a function of the number of active duck farms and the mean distance between active poultry farms (km). The black lines indicate increases in *R*_e_ values by increments of 1.

### Density and proximity thresholds limiting HPAI transmission between farms

From the associated predictors, we focused on those that can inform management actions, specifically the number of active duck farms and their proximity from each other. Using the coefficients of these two predictors, we estimated the effective reproduction number (*R*_e_) between farms (**Figure 3B**) and identified thresholds that ensure *R*_e_ < 1. Our results indicate that in areas with a high number of active duck farms, greater distances between poultry farms are necessary to reduce transmission risks.

## Discussion

Our study presents a comprehensive phylodynamic analysis of the transmission dynamics of the HPAI H5N8 (clade 2.3.4.4b) virus in Southwestern France in 2020-21. The virus lineage was likely introduced by migratory wild birds within a short time frame in early December, consistent with migratory routes descending during autumn ^21^. Following its introduction, the virus was maintained in duck flocks in Landes before spreading to neighbouring departments. The transmission dynamics peaked in mid-December 2020, with an estimated velocity of 27.8km/week and a peak effective reproduction number (*R*_e_) of approximately 3.8. Notably, this *R*_e_ value was higher than the peak observed during the 2016-17 wave which was estimated at around 1.5^22^. This suggests that the virus was able to spread beyond the standard 3- and 10-km control and surveillance zones. This widespread transmission was likely exacerbated by a prolonged silent phase: infected poultry farms were estimated to remain infectious (from infection to suspicion) for around 9 days, consistent with previous estimations ^23^. This highlights a critical window of undetected viral spread, during which infected poultry, particularly ducks, can shed high viral loads into the environment.

The subsequent decline in *R*_e_ and velocity after late December coincided with extensive preventive culling, which targeted all poultry species within a 1-km radius of infected farms and all ducks and other outdoor poultry species within a 3 or 5-km radius of infected farms, likely contributing to the observed decrease in transmission ^24^.

We identified key factors driving the spread of the virus between poultry farms. Our results indicate that, following the introduction of the viral lineage in Southwestern France, subsequent poultry farm outbreaks were primarily driven by high duck farm densities in the region. This is supported by the higher susceptibility and infectiousness of ducks to circulating HPAI H5N8 compared to chickens ^22,25,26^. This effect was even more pronounced when poultry farms were located closer to each other. However, the underlying mechanisms and their contributions to transmission in high-density areas remain unclear, as detailed data is lacking to truly test their contributions. Previous studies have shown high virus contamination in the vicinity of poultry farms, up to several hundred meters ^27^. Given the high duck farm density in the region, this suggests that contaminated dust or feathers could reach neighbouring farms. Even after cleaning and disinfection, virus particles have been detected in trucks transporting infected ducks^28^, further suggesting that local spread culd occur along transport routes. Infection by proximity may also be linked to human activity (e.g. farm workers, veterinarians, or equipment moving between farms), especially given the high connectivity between farms, or to local wildlife (e.g. commensal birds living near farms ^29^) which could potentially play a bridging role. Moreover, the rapid increase in outbreaks, characterized by high numbers and short time intervals between suspicions (on average, half a day), may have overwhelmed the capacity for culling infected flocks and preventive depopulation, despite massive efforts. This resulted in delays between suspicion and culling, likely contributing to the persistence of environmental virus contamination. All of these factors underscore the territorial dimension of the risk of transmission, where virus spread is likely driven not by a single factor but by a combination of factors, particularly in densely connected regions.

Heavy rainfall in winter 2020-21 caused prolonged water retention in outdoor areas, creating ideal conditions for virus survival ^24^. While this raised concerns about water runoff facilitating virus spread between farms, our analysis using rainfall as a proxy did not show a significant association with transmission. It is however possible that our rainfall data did not fully capture the complexity of water runoff dynamics, warranting further investigation. In contrast, the percentage of water surface areas, used as proxy of wild bird densities, was positively associated with between-farm transmission, potentially reflecting the involvement of wild birds as bridge hosts ^30^. However, only a small number of wild birds, mostly large species like swans and geese, were reported infected in France in 2020-21 ^31^. This raises the possibility that other species, such as smaller birds that are less likely to be found if dead or species not targeted by the surveillance system ^32^, could play a role in the virus transmission between farms. Incorporating more specific wild bird species densities and movement data could enhance future outbreak predictions ^33,34^. Other factors influencing disease spread could not be assessed in this study. These included personnel and equipment movement, which should be further investigated. For now, raising awareness among farm workers, veterinarians and technicians about biosecurity measures when moving between farms is essential.

Although poultry movements were not included here, previous studies in France have shown a limited impact ^35,36^. Additional data on flock size and farming practices (e.g., outdoor versus indoor systems) were unavailable but remain critical for future research. Notably, in late 2020, many exemptions were granted to compulsory indoor housing requirements, due to logistical and economic challenges. However, efforts are still ongoing to better account for species and production-specific constraints while adapting measures to the level of risk, reflecting the continued search for practical and collaborative solutions with farmers.

Our study provides evidence-based thresholds for farm density and proximity to prevent virus spread within the country, complementing existing farm-level biosecurity measures. These findings area particularly relevant for regions with high poultry farm densities and outdoor farming practices, which inherently increase the risk of sustained transmission once the virus is introduced. While vaccination has been implemented and has contributed to a lower number of outbreaks since late 2023, other measures remain necessary to ensure long-term control. To mitigate future outbreaks, our study offers actionable guidance to support regional biosecurity and rethink the spatial distribution and organisation of poultry farms ^11^. Moving forward, implementing these thresholds may pose challenges, but short-term solutions like increasing downtime between production cycles or limiting poultry placements during high-risk periods, as successfully implemented by the poultry industry-driven plan (“Plan Adour”), could help, providing adequate financial compensation. In the long term, structural changes like promoting new production sites in low-density areas or enforcing regulatory frameworks for minimal farm distances such as Italy’s recent Ministry of Health decree ^37^, may be necessary. However, these strategies warrant further investigation to evaluate their socio-economic impacts and feasibility.

## Methods

### Selection and curation of viral sequences

A total of 468 poultry farm outbreaks of HPAI H5N8 were reported from December 2, 2020 to March 20, 2021 in France (**Figure 1**). Among these outbreaks, 83.1% (381/468) occurred in duck farms located in South-western France. During this period, the National Reference Laboratory for Avian Influenza (LNR IA, Anses-Ploufragan-Plouzané-Niort) generated a total of 432 sequences of HA segment ^31^. To identify introduction events into France, we merged and aligned these sequences with those available on GISAID (https://gisaid.org) in Europe from September 7, 2020 to May 1, 2021 (n = 613 sequences) using MAFFT v7.49 ^38^. After subsampling to remove identical sequences with the same sampling date, species and location, we inferred a maximum likelihood phylogenetic tree using the HKY+Γ_4_ nucleotide substitution model using RAxML-NG ^5339^.

### Reconstructing the spatio-temporal dispersal history of viral lineages

To infer spatially and temporally referenced phylogenetic trees, we performed a continuous phylogeographic analysis using BEAST v1.10.4 ^40^ and the BEAGLE library ^41^ to improve computational performance. The tree prior was specified as a skygrid coalescent model ^42^. The model was coupled with a HKY+Γ4 nucleotide substitution model ^43^ and an uncorrelated relaxed molecular clock following a lognormal prior distribution ^44^. The relaxed random walk (RRW) diffusion model ^45^ was used to perform the continuous phylogeographic reconstruction using the geographical coordinates of the sequences. Analyses were run for 100 million steps across three independent Markov chains, with states sampled every 100,000 steps. Convergence and mixing properties were assessed using Tracer v1.7.1 ^46^. After discarding 10% of sampled posterior trees as burn-in, the maximum clade credibility (MCC) tree was generated using TreeAnnotator v1.10.4 ^40^ and annotated using the “ggtree” package in R v4.2.2 ^47^. The spatiotemporal information embedded within the posterior trees was extracted and visualized using the “seraphim” package ^48^.

### Quantifying the between-farm transmission dynamics of viral lineages

To infer the between-farm effective reproductive number (*R*_e_) over time, we performed a phylodynamic analysis using BEAST v2.7.5 ^49^ and the BEAGLE library ^41^. This parameter was defined as the average number of secondary poultry farms infected by one infectious poultry farm. The tree prior was specified as a single-type birth-death-sampling model using the “BDMM-Prime” package (https://github.com/tgvaughan/BDMM-Prime) ^50,51^. The model assumes a constant *R*_*e*_ before December 12, 2020, after February 27, 2021 and infers a weekly *R*_*e*_ estimates between those dates. We assumed that no poultry farm outbreaks went unobserved, considering the severity of the clinical signs, especially in ducks, and the recurrent nature of HPAI waves, raising farmers’ awareness. Thus, the sampling proportion was informed by the number of sequences over the number of reported poultry farm outbreaks. The infectious duration of farm units (e.g. the time from farm infection to sampling/culling) was approximated by a prior distribution around a median of 7 days to match previous epidemiological estimates ^22,23^. It was assumed that once sampled, a given farm could not be infected and sampled again; since infected poultry farms were subject to culling following the confirmation of infection. The origin of the epidemic was approximated by a prior distribution around 7 days prior to the first reported outbreak to align with previous epidemiological estimates^22,23^. The model was coupled with a HKY+Γ4 nucleotide substitution model ^43^ and a lognormally distributed uncorrelated relaxed molecular clock ^44^. More details about the prior values and distributions of the model parameters are described in **Table S1**. Analyses were run for 2 billion steps across three independent Markov chains, with states sampled every 200,000 steps. Convergence and mixing properties were assessed using Tracer v1.7.2 ^46^. After discarding 10% of sampled posterior trees as burn-in, weekly *R*_e_ estimates were plotted over time and compared to those previously estimated based on epidemiological data only ^8^ using the “EpiEstim” package ^52^ in R v4.2.2 ^61^.

### Testing the impact of drivers on the between-farm transmission of viral lineages

To investigate the impact of specific predictors on the between-farm effective reproductive number (*R*_e_), the single-type birth-death-sampling model was extended with a generalized linear model (GLM) using the GLMPrior package (https://github.com/cecivale/GLMPrior). In this parametrization, the weekly *R*_e_ parameter acts as the outcome to a log-linear function of weekly mean predictor values. Predictors were: (i) the *number of active duck farms*, which refers to the weekly average number of duck farms with poultry present (excluding those that were empty due to sanitary procedures between production cycles, or due to culling measures, if they became infected or preventively culled) within a 10 km radius of outbreaks reported that week and in the 7 days before they were reported. The 7-day period was based on the average time it takes to detect infected farms ^22,23^ and the 10 km distance aligned with the radius of the regulatory control zones in where specific measures are implemented to prevent further disease spread, such as movement restrictions, enhanced surveillance, and biosecurity protocols. Sensitivity analyses also considered a 12-day period (source: DGAl). (ii) the *number of active chicken farms*, which is similar to the active duck farms, e.g., the weekly average number of chicken farms with poultry present within a space-time window of 10 km and 7 days prior to outbreak reporting (source: DGAl). (iii) the *mean distances between active poultry farms*, which measures the weekly average distance between all poultry farms (both duck and chicken) that were active within a space-time window of 10 km and 7 days prior to outbreak reporting (source: DGAl). (iv) the *mean percentage of water surface areas*, which represents the weekly average percentage of areas covered by water bodies (such as ponds, rivers) within a 10 km radius of outbreaks, serving as an indicator of the presence of wild aquatic birds (source: Système d’Information sur l’Eau, https://www.data.gouv.fr/fr/datasets/cours-deau-metropole-2017-bd-carthage/). (v) the *mean rainfall*, which refers to the weekly average amount of rainfall in millimeters reported by the closest weather station to each outbreak and in the 7 days prior to outbreak reporting (source: Météo-France https://meteo.data.gouv.fr/). Temporal distribution of the weekly mean predictor values can be found in **Figures S2** and **S3**. Following collinearity assessment among predictors, all predictors were kept given that all absolute values of the Pearson correlation were below 0.7 ^53^.

For each predictor, the GLM parameterization also includes a regression coefficient which quantifies the (log) contribution of the predictors and a binary indicator variable indicating whether the predictor is included in the model. Averaging that binary variable across all steps quantifies the probability of each predictor to be included in the model. To reduce the effect of different predictors’ magnitude, all non-binary predictors were log-transformed and standardized before inclusion in the GLM. Bayes Factors (BF) were used to determine the contribution of each predictor in the GLM ^12,54,55^. BF were calculated for each predictor to quantify which of the posterior and prior inclusion probabilities of the given predictor in the model is more likely. A cutoff of 3.2, 10 and 100 was used to indicate substantial, strong and decisive contribution of a predictor in the GLM, respectively ^54^, meaning that its posterior inclusion probability in the model was 3.2-, 10- or 100-fold more likely than its prior inclusion probability (0.50), respectively. To provide practical insights, predictions of the between-farm effective reproductive number (*R*_e_) were made based on key predictors showing potential for actionable interventions, using their posterior coefficients when they were the only predictors included in the model.

## Data availability

All HPAI genetic sequences used in this study are available on the GISAID database (http://www.gisaid.org) and upon request from the French national reference laboratory for avian influenza viruses (https://www.anses.fr). The accession numbers in GISAID are available from https://github.com/ClaireGuinat/h5n8_glm_phylo.

Poultry farm structure and outbreak management data are available upon request from the French Ministry of Agriculture and Food (https://agriculture.gouv.fr/french-ministry-agriculture-and-food). Climatic and landscape data are accessible via Météo-France (https://meteo.data.gouv.fr/) and Système d’Information sur l’Eau (https://www.data.gouv.fr/fr/datasets/cours-deau-metropole-2017-bd-carthage/) databases.

## Code availability

The XML files used to perform the phylogeographic and phylodynamic analyses are available from https://github.com/ClaireGuinat/h5n8_glm_phylo.

## Acknowledgements

We thank the veterinarians and farmers involed in samples collection, the local official laboratories for initial samples screening, the local and central veterinary services (Directions Départementales en charge de la Protection des Populations, Direction Générale de l’Alimentation) involved in the HPAI outbreak management and the National Reference Laboratory for Avian Influenza viruses (LNR AI ANSES-Ploufragan-Plouzané-Niort) for the sequencing process. The authors gratefully acknowledge the contributions from other laboratories to GISAID.

## Funding statement

This work was financially supported by the FEDER/Région Occitanie Recherche et Sociétés 2018—AI-TRACK and the PREDYT project (Fonds pour la Recherche sur l’Influenza Aviaire). This study was also supported by the “Chair for Avian Biosecurity”, hosted by the National Veterinary School of Toulouse and funded by the Direction Générale de l’Alimentation, France.

## Author contributions

Study design: C.G., D.C., M.C.P., T.V. Data resources: F-X.B., B.G., L.F. Data curation: L.F., Data analysis: C.G., S.L. Package resources: C.V.A., Analysis guidance: C.V.A., T.S., L.D.P., Manuscript preparation: C.G. Review and approval of final manuscript: all authors

## Competing interests

The authors declare no competing interests.

## Notes

### Competing Interest Statement

The authors have declared no competing interest.

## References

1. Xie, R. et al. The episodic resurgence of highly pathogenic avian influenza H5 virus. Nature 622, 810–817 (2023).

2. Puryear, W. B. & Runstadler, J. A. High-pathogenicity avian influenza in wildlife: a changing disease dynamic that is expanding in wild birds and having an increasing impact on a growing number of mammals. Journal of the American Veterinary Medical Association 1, 1–9 (2024).

3. Plaza, P. I., Gamarra-Toledo, V., Euguí, J. R. & Lambertucci, S. A. Recent Changes in Patterns of Mammal Infection with Highly Pathogenic Avian Influenza A (H5N1) Virus Worldwide. Emerging Infectious Diseases 30, 444 (2024).

4. Bi, Y., Yang, J., Wang, L., Ran, L. & Gao, G. F. Ecology and evolution of avian influenza viruses. Current Biology 34, R716–R721 (2024).

5. European Food Safety Authority (EFSA) et al. Drivers for a pandemic due to avian influenza and options for One Health mitigation measures. EFSA Journal 22, e8735 (2024).

6. Koopmans, M. P. G. et al. The panzootic spread of highly pathogenic avian influenza H5N1 sublineage 2.3.4.4b: a critical appraisal of One Health preparedness and prevention. The Lancet Infectious Diseases doi:10.1016/S1473-3099(24)00438-9.

7. Guinat, C. et al. Spatio-temporal patterns of highly pathogenic avian influenza virus subtype H5N8 spread, France, 2016 to 2017. Eurosurveillance 23, 1700791 (2018).

8. Lambert, S. et al. Two major epidemics of highly pathogenic avian influenza virus H5N8 and H5N1 in domestic poultry in France, 2020–2022. Transboundary and Emerging Diseases (2022).

9. Scoizec, A. et al. New Patterns for Highly Pathogenic Avian Influenza and Adjustment of Prevention, Control and Surveillance Strategies: The Example of France. Viruses 16, 101 (2024).

10. Sidik, S. M. How to stop the bird flu outbreak becoming a pandemic. (2023).

11. Vergne, T. et al. Highly pathogenic avian influenza management policy in domestic poultry: from reacting to preventing. Eurosurveillance 29, 2400266 (2024).

12. Lemey, P. et al. Unifying viral genetics and human transportation data to predict the global transmission dynamics of human influenza H3N2. PloS pathog 10, e1003932 (2014).

13. Baele, G., Suchard, M. A., Rambaut, A. & Lemey, P. Emerging concepts of data integration in pathogen phylodynamics. Systematic biology 66, e47–e65 (2017).

14. Hicks, J. T. et al. Agricultural and geographic factors shaped the North American 2015 highly pathogenic avian influenza H5N2 outbreak. PLoS Pathog 16, e1007857 (2020).

15. Gass Jr, J. D. et al. Ecogeographic drivers of the spatial spread of highly pathogenic avian influenza outbreaks in Europe and the United States, 2016–Early 2022. International Journal of Environmental Research and Public Health 20, 6030 (2023).

16. Bui, C. M., Adam, D. C., Njoto, E., Scotch, M. & MacIntyre, C. R. Characterising routes of H5N1 and H7N9 spread in China using Bayesian phylogeographical analysis. Emerging microbes & infections 7, 1–8 (2018).

17. Yang, J., Müller, N. F., Bouckaert, R., Xu, B. & Drummond, A. J. Bayesian phylodynamics of avian influenza A virus H9N2 in Asia with time-dependent predictors of migration. PLoS Comput Biol 15, e1007189 (2019).

18. Yang, Q. et al. Assessing the role of live poultry trade in community-structured transmission of avian influenza in China. Proceedings of the National Academy of Sciences 117, 5949–5954 (2020).

19. Carnegie, L. et al. H9N2 avian influenza virus dispersal along Bangladeshi poultry trading networks. Virus Evolution 9, vead014 (2023).

20. Lu, L., Brown, A. J. L. & Lycett, S. J. Quantifying predictors for the spatial diffusion of avian influenza virus in China. BMC evolutionary biology 17, 1–14 (2017).

21. Guillemain, M., Plaquin, B., Caizergues, A., Bacon, L. & De Wiele, A. V. La migration des anatidés: patron général, évolutions, et conséquences épidémiologiques. Bull. épidemiologique, santé Anim. Aliment (2021).

22. Andronico, A. et al. Highly pathogenic avian influenza H5N8 in south-west France 2016–2017: A modeling study of control strategies. Epidemics 28, 100340 (2019).

23. Vergne, T. et al. Inferring within□flock transmission dynamics of highly pathogenic avian influenza H5N8 virus in France, 2020. Transboundary and emerging diseases 68, 3151–3155 (2021).

24. CES Saba. AVIS de l’Agence Nationale de Sécurité Sanitaire de l’alimentation, de l’environnement et Du Travail Relatif à Un Retour d’expérience Sur La Crise Influenza Aviaire Hautement Pathogène 2020–2021 (1ère Partie). [Available at: Https://Www.Anses.Fr/Fr/System/Files/SABA2021SA0022-2.Pdf (Accessed in September 2024)]. (2021).

25. Bertran, K. et al. Lack of chicken adaptation of newly emergent Eurasian H5N8 and reassortant H5N2 high pathogenicity avian influenza viruses in the U.S. is consistent with restricted poultry outbreaks in the Pacific flyway during 2014–2015. Virology 494, 190–197 (2016).

26. Grund, C. et al. A novel European H5N8 influenza A virus has increased virulence in ducks but low zoonotic potential. EMERGING MICROBES & INFECTIONS 7, (2018).

27. Scoizec, A. et al. Airborne Detection of H5N8 Highly Pathogenic Avian Influenza Virus Genome in Poultry Farms, France. Front. Vet. Sci. 5, (2018).

28. Huneau-Salaün, A., Scoizec, A., Thomas, R. & Le Bouquin, S. Cleaning and disinfection of crates and trucks used for duck transport: field observations during the H5N8 avian influenza outbreaks in France in 2017. Poultry Science 99, 2931–2936 (2020).

29. Gall-Ladevèze, L. et al. Quantification and characterisation of commensal wild birds and their interactions with domestic ducks on a free-range farm in southwest France. Scientific Reports 12, 1–13 (2022).

30. Caron, A., Cappelle, J., Cumming, G. S., de Garine-Wichatitsky, M. & Gaidet, N. Bridge hosts, a missing link for disease ecology in multi-host systems. Veterinary Research 46, 1–11 (2015).

31. Briand, F. et al. Multiple independent introductions of highly pathogenic avian influenza H5 viruses during the 2020–2021 epizootic in France. Transboundary and emerging diseases 69, 4028–4033 (2022).

32. Decors, A. et al. Le réseau Sagir: un outil de vigilance vis-à-vis des agents pathogènes exotiques. Bull. Épidémiologique Santé Anim.-Aliment 66, 35–39 (2014).

33. Schreuder, J. et al. Wild bird densities and landscape variables predict spatial patterns in HPAI outbreak risk across The Netherlands. Pathogens 11, 549 (2022).

34. Wu, H.-D. I. et al. Integrating Citizen Scientist Data into the Surveillance System for Avian Influenza Virus, Taiwan. Emerging Infectious Diseases 29, 45 (2023).

35. Guinat, C. et al. Role of Live-Duck Movement Networks in Transmission of Avian Influenza, France, 2016–2017. Emerg Infect Dis 26, 472–480 (2020).

36. Bauzile, B. et al. Unravelling direct and indirect contact patterns between duck farms in France and their association with the 2016–2017 epidemic of Highly Pathogenic Avian Influenza (H5N8). Preventive Veterinary Medicine 198, 105548 (2022).

37. Decreto Ministeriale. Modalita’ Applicative Delle Misure Di Biosicurezza Negli Allevamenti Avicoli. (23A03711) (GU Serie Generale n.151 Del 30–06-2023) [Available at: Https://Www.Gazzettaufficiale.It/Atto/Serie_generale/caricaDettaglioAtto/Originario?Atto.dataPubblicazioneGazzetta=2023-06-30&atto.codiceRedazionale=23A03711&elenco30giorni=false x(Accessed in September 2024)]. (2023).

38. Katoh, K. & Standley, D. M. MAFFT Multiple Sequence Alignment Software Version 7: Improvements in Performance and Usability. Molecular Biology and Evolution 30, 772–780 (2013).

39. Kozlov, A. M., Darriba, D., Flouri, T., Morel, B. & Stamatakis, A. RAxML-NG: a fast, scalable and user-friendly tool for maximum likelihood phylogenetic inference. Bioinformatics 35, 4453–4455 (2019).

40. Drummond, A. J., Suchard, M. A., Xie, D. & Rambaut, A. Bayesian phylogenetics with BEAUti and the BEAST 1.7. Mol Biol Evol 29, 1969–1973 (2012).

41. Ayres, D. L. et al. BEAGLE: an application programming interface and high-performance computing library for statistical phylogenetics. Systematic biology 61, 170–173 (2012).

42. Hill, V. & Baele, G. Bayesian estimation of past population dynamics in BEAST 1.10 using the Skygrid coalescent model. Molecular biology and evolution 36, 2620–2628 (2019).

43. Shapiro, B., Rambaut, A. & Drummond, A. J. Choosing appropriate substitution models for the phylogenetic analysis of protein-coding sequences. Molecular biology and evolution 23, 7–9 (2006).

44. Drummond, A. J., Ho, S. Y. W., Phillips, M. J. & Rambaut, A. Relaxed phylogenetics and dating with confidence. PLoS Biol 4, e88 (2006).

45. Lemey, P., Rambaut, A., Welch, J. J. & Suchard, M. A. Phylogeography takes a relaxed random walk in continuous space and time. Molecular biology and evolution 27, 1877–1885 (2010).

46. Rambaut, A., Drummond, A. J., Xie, D., Baele, G. & Suchard, M. A. Posterior summarization in Bayesian phylogenetics using Tracer 1.7. Systematic biology 67, 901 (2018).

47. R Core Team, R. R: A language and environment for statistical computing. (2013).

48. Dellicour, S., Rose, R., Faria, N. R., Lemey, P. & Pybus, O. G. SERAPHIM: studying environmental rasters and phylogenetically informed movements. Bioinformatics 32, 3204–3206 (2016).

49. Bouckaert, R. et al. BEAST 2: a software platform for Bayesian evolutionary analysis. PLoS Comput Biol 10, e1003537 (2014).

50. Scire, J., Barido-Sottani, J., Kühnert, D., Vaughan, T. G. & Stadler, T. Improved multi-type birth-death phylodynamic inference in BEAST 2. BioRxiv (2020).

51. Kühnert, D., Stadler, T., Vaughan, T. G. & Drummond, A. J. Phylodynamics with migration: a computational framework to quantify population structure from genomic data. Molecular biology and evolution 33, 2102–2116 (2016).

52. Cori, A., Ferguson, N. M., Fraser, C. & Cauchemez, S. A new framework and software to estimate time-varying reproduction numbers during epidemics. American journal of epidemiology 178, 1505–1512 (2013).

53. Dohoo, I. R., Martin, W. & Stryhn, H. E. Veterinary Epidemiologic Research. (2003).

54. Kass, R. E. & Raftery, A. E. Bayes Factors. Journal of the American Statistical Association 90, 773–795 (1995).

55. Magee, D., Beard, R., Suchard, M. A., Lemey, P. & Scotch, M. Combining phylogeography and spatial epidemiology to uncover predictors of H5N1 influenza A virus diffusion. Archives of virology 160, 215–224 (2015).

