## Supplementary files for "Highly pathogenic avian influenza management in high-density poultry farming areas"

**Supplementary Figures**

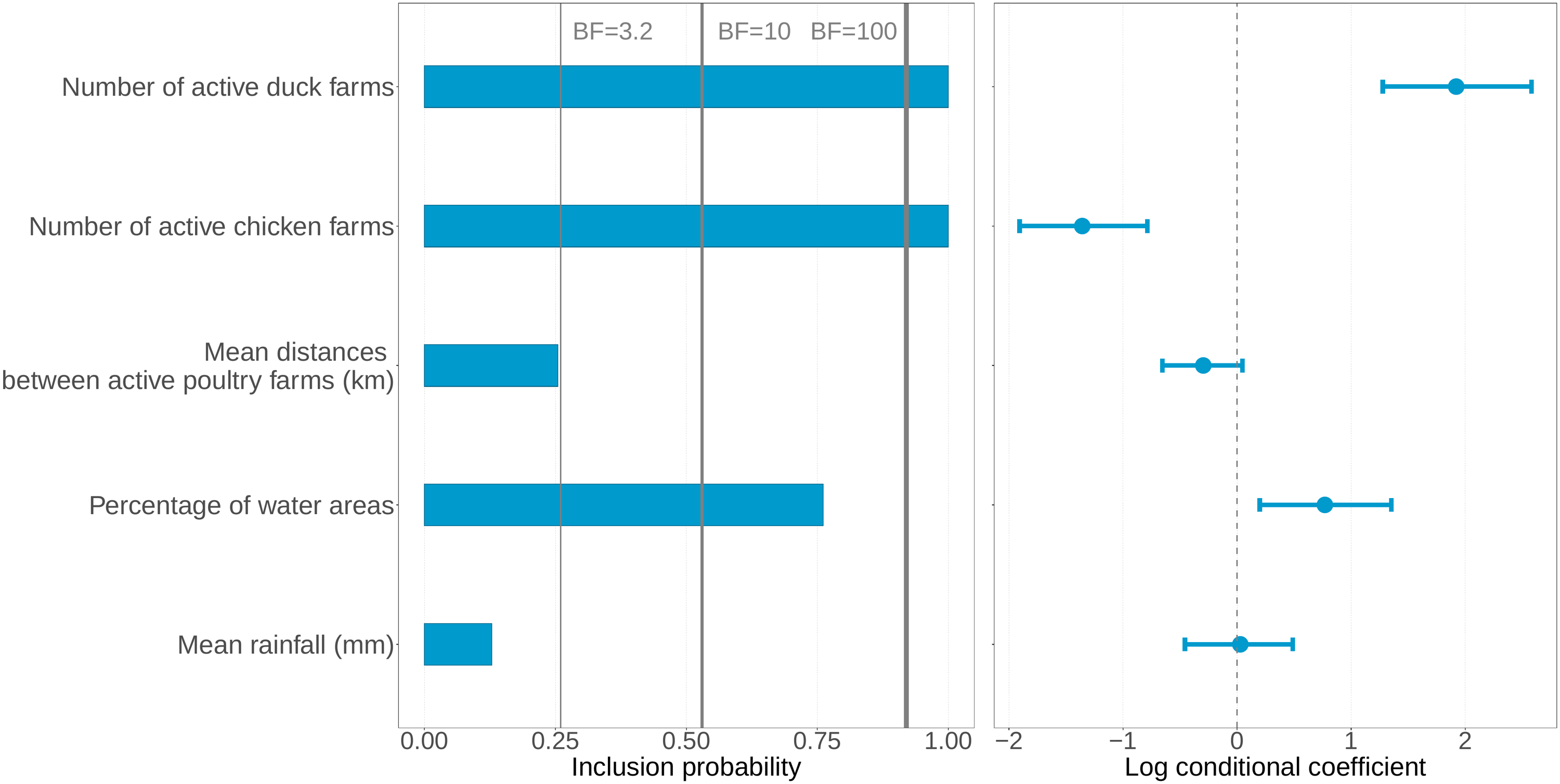

**Figure S1.** Drivers of HPAI H5N8 clade 2.3.4.4b transmission between farms in France 2020-21 using a 12-day period as sensitivity analysis

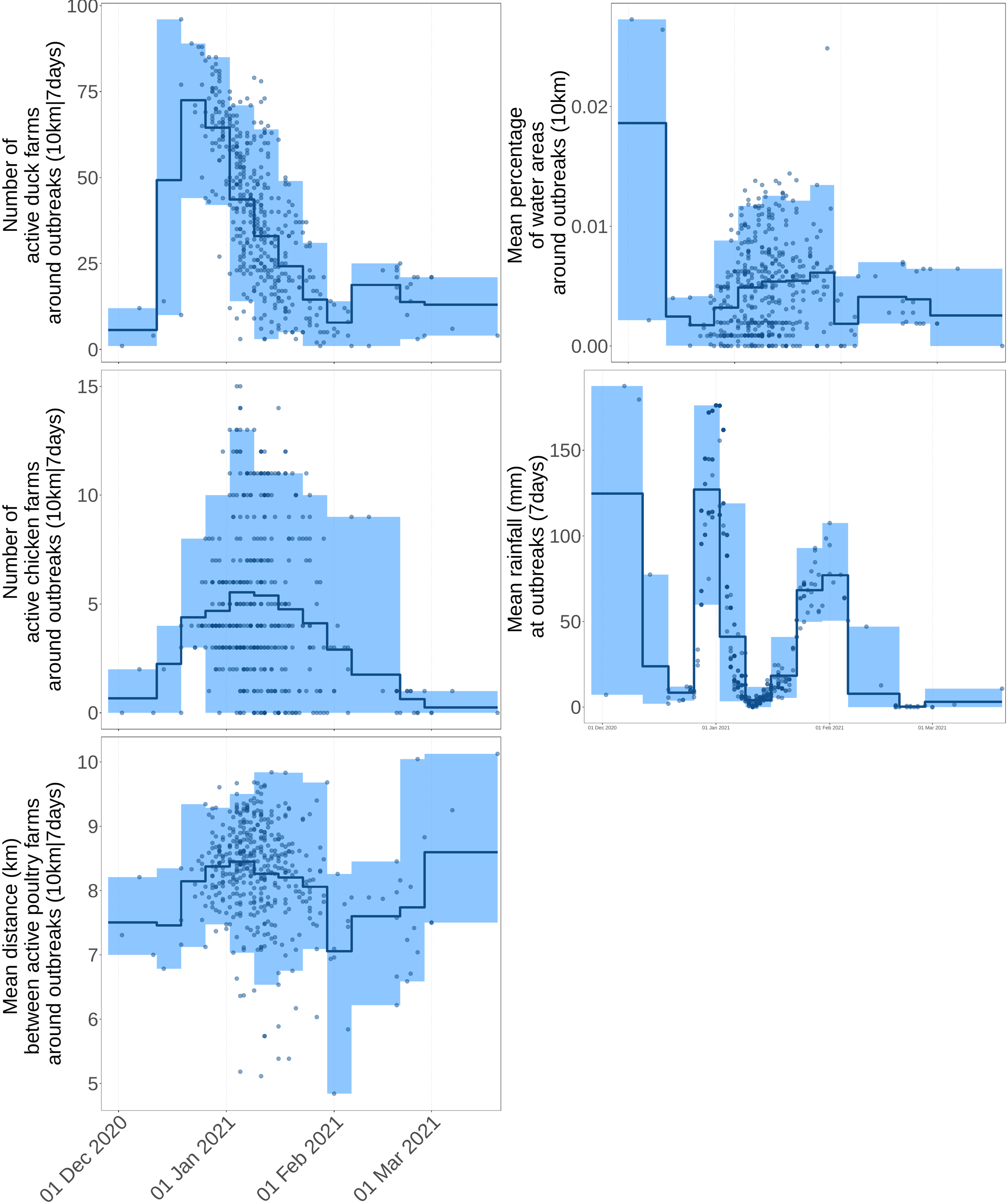

**Figure S2.** Temporal distribution of the predictors tested on the weekly between-farm effective reproductive number. Each blue dot represent the value of the predictor calculated for a given outbreak reported on the date displayed on the x-axis within a 10 km radius of outbreaks reported that week and in the 7 days before they were reported. The horizontal blue line represents the weekly average value of the predictor and the blue shaded area the highest density interval.
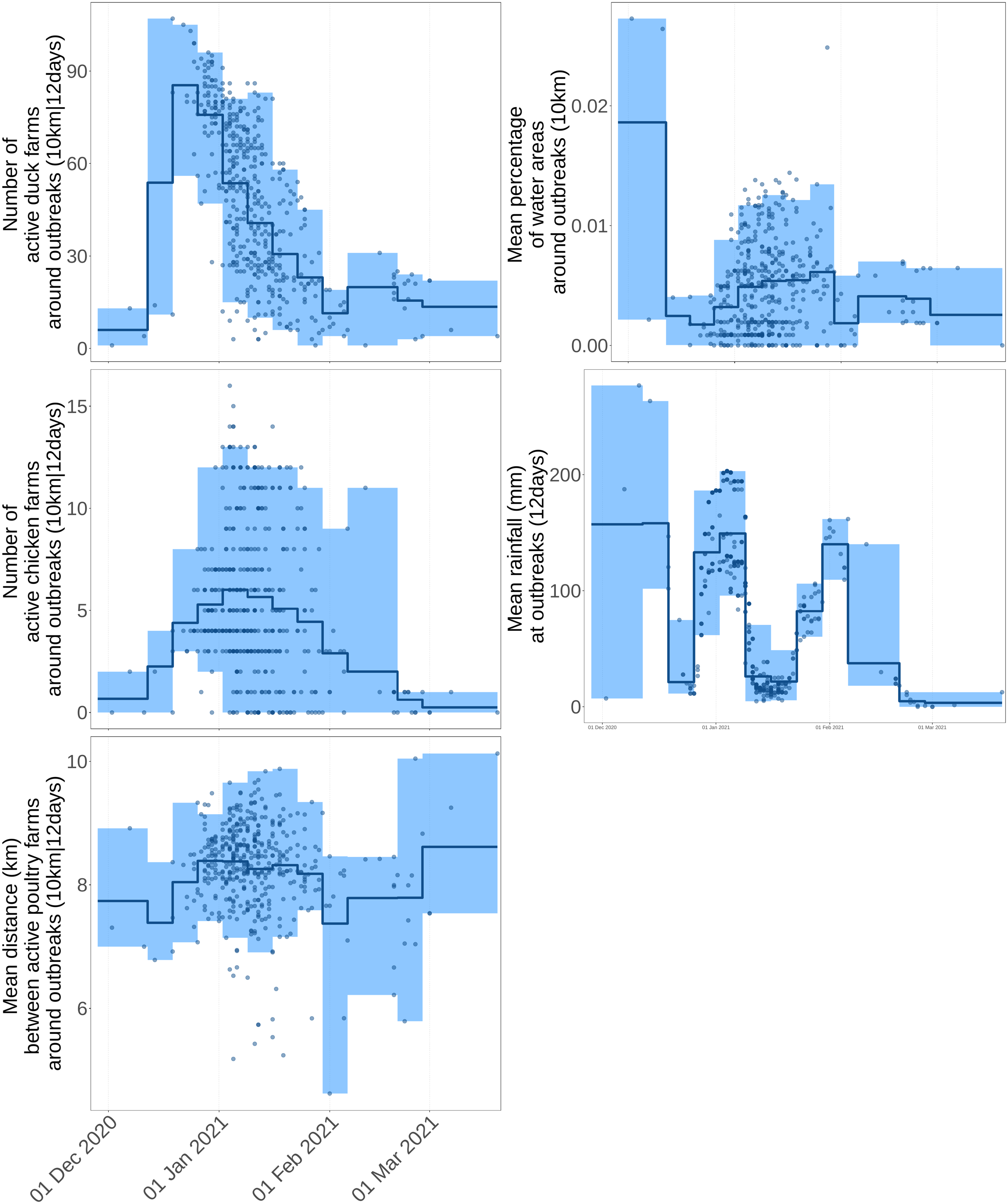

**Figure S3.** Temporal distribution of the predictors tested on the weekly between-farm effective reproductive number. Each blue dot represent the value of the predictor calculated for a given outbreak reported on the date displayed on the x-axis within a 10 km radius of outbreaks reported that week and in the 12 days before they were reported. The horizontal blue line represents the weekly average value of the predictor and the blue shaded area the highest density interval.

**Supplementary Tables**

**Table S1.** Prior values and distributions used in the single-type birth-death model

| Parameter | Prior | Rationale |
| --- | --- | --- |
| Nucleotide substitution model | HKY + Γ_4_ | Unequal transition/transversion rates, unequal base frequencies, rate heterogeneity among sites with four categories |
| Molecular clock rate | Mean rate across branches fixed to 0.001 | 0.001 substitution/site/year |
| Become uninfectious rate | Lognormal(52,0.6) | Correspond to a median of 43.4/year [95% IQR: 13.4-141], e.g. a median of 7 days [95% IQR: 2 - 27] |
| Sampling proportion | Fixed to 0 before 28 November 2020  Fixed to 0.8211 after 28 November 2020 | 0 correspond to no sequences collected before the first reported outbreak  0.8211 corresponds to 381 sequences collected over 464 reported outbreaks |
| Time of origin | Lognormal(-1.4,0.05) | Correspond to a median of 19 December 2020 [95% IQR: 10 December 2020 – 28 December 2021] |
| Regression coefficient | Normal(0,2) | Correspond to a median of 0 [95% IQR: -3.92 – 3.92] |
| Indicator variable | Binary: 0 or 1 | 0.5 prior inclusion probability for each predictor |
